## Supplementary material for "Diffusion MRI Anisotropy in the Cerebral Cortex is Determined by Unmyelinated Tissue Features": Supplemetary Informaton (SI) Figures

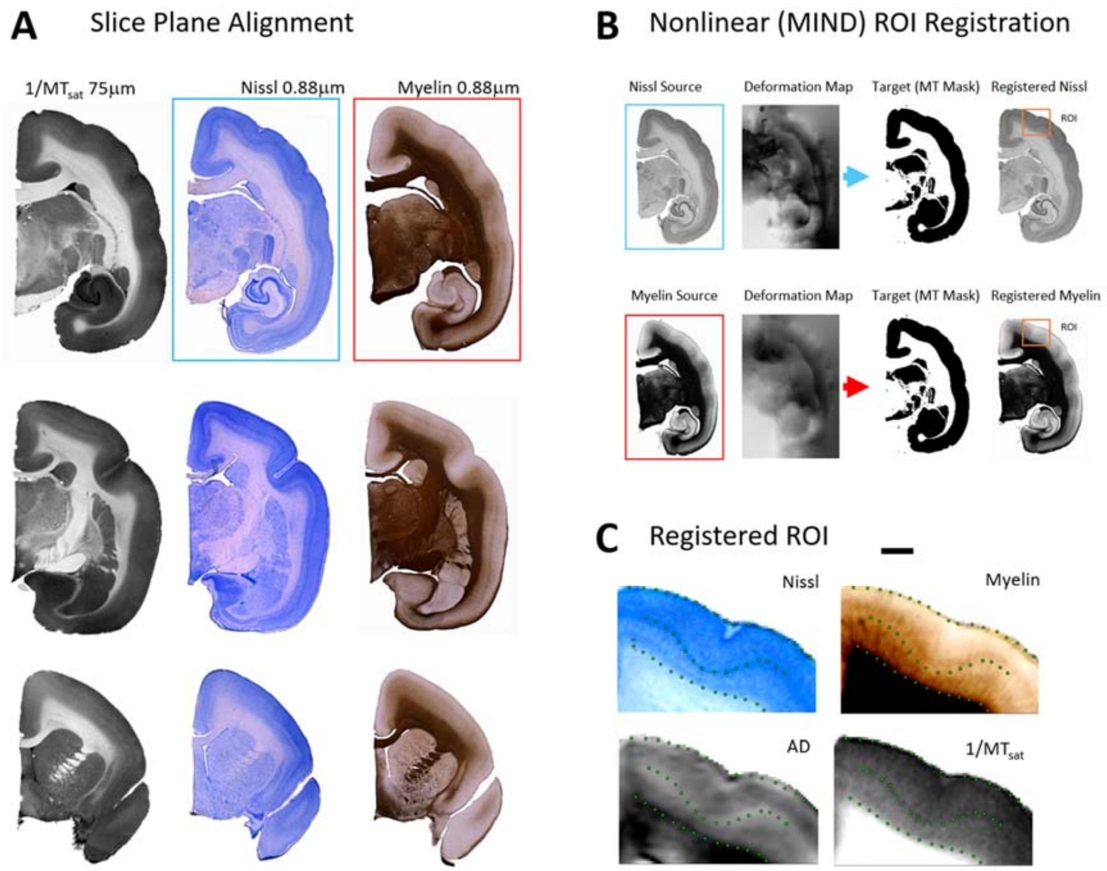


*SI Figure 1 Co-registration of histology to MRI involved a rigid body rotation of MRI data to match the histology slice plane, followed by non-linear correction for histology shrinkage and deformation using MIND **a)** Examples of Nissl, Myelin and MT at slices, after rotation to slice plane alignment. **b)** Non-linear registration to featureless masks of MT data. **c)** registered data, points show granular layer 4, white gray boundary and pial boundary*

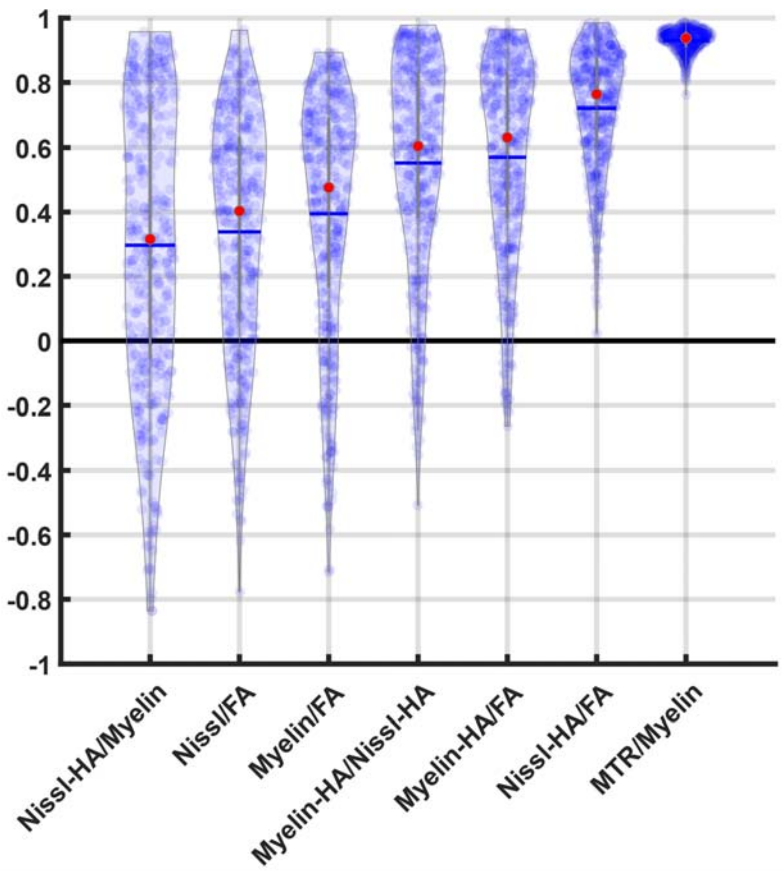


*SI Figure 2 Distributions of correlation coefficients between MRI and histological variables between MRI and histological variables*

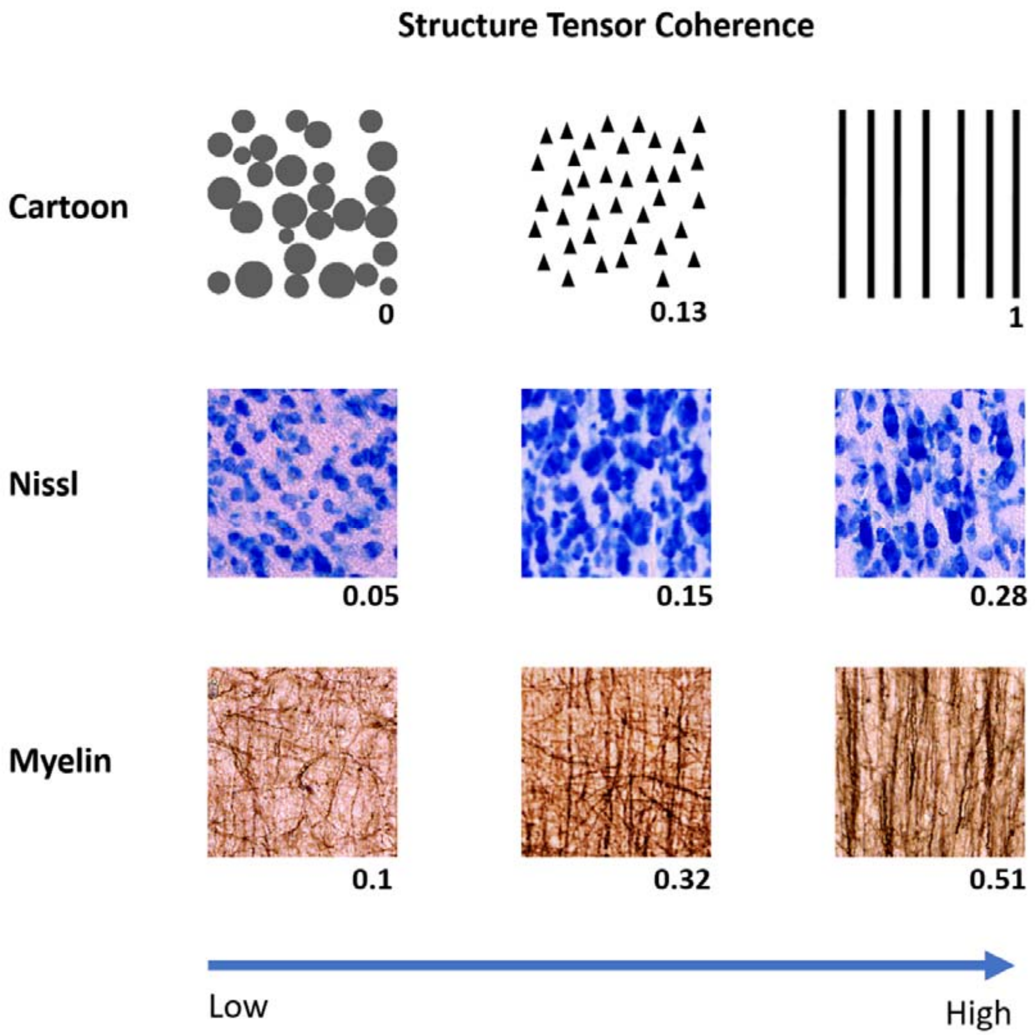


***SI Figure 3*** *Weighted ST anisotropy values for 75um areas of histology and cartoon data (see Methods)*

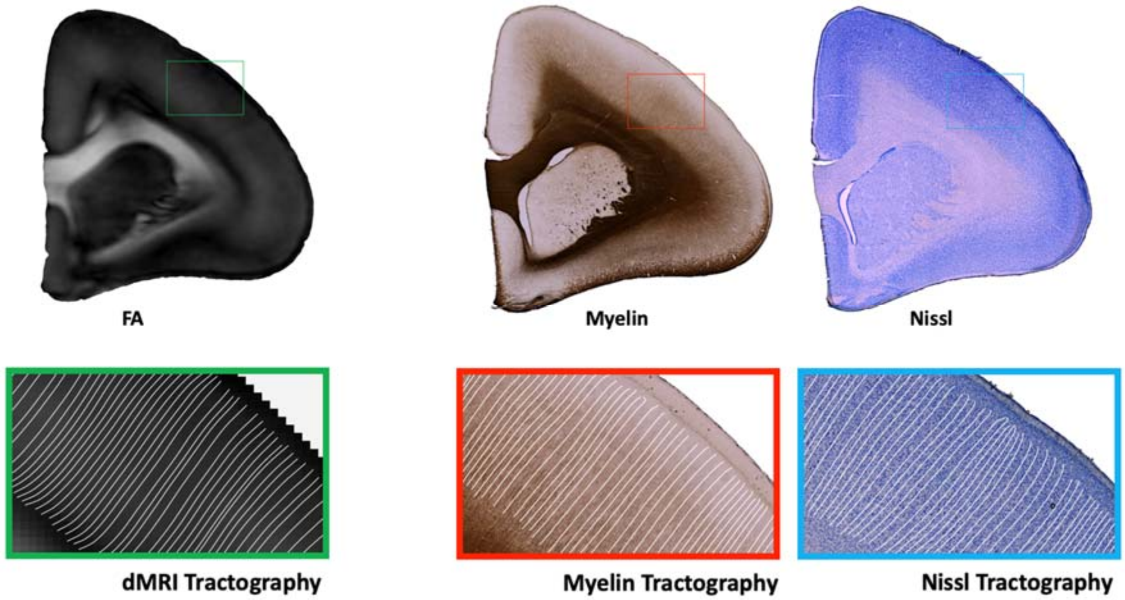


***SI Figure 4*** *Examples of dMRI tractography and tractography through the vector field defined by the histology weighted structure tensors (see* ***Methods****). Left: tractography through the 2D projected principle eigenvector of the diffusion tensor. Middle tractography through the myelin structure tensor. Right tractography through the Nissl structure tensor*

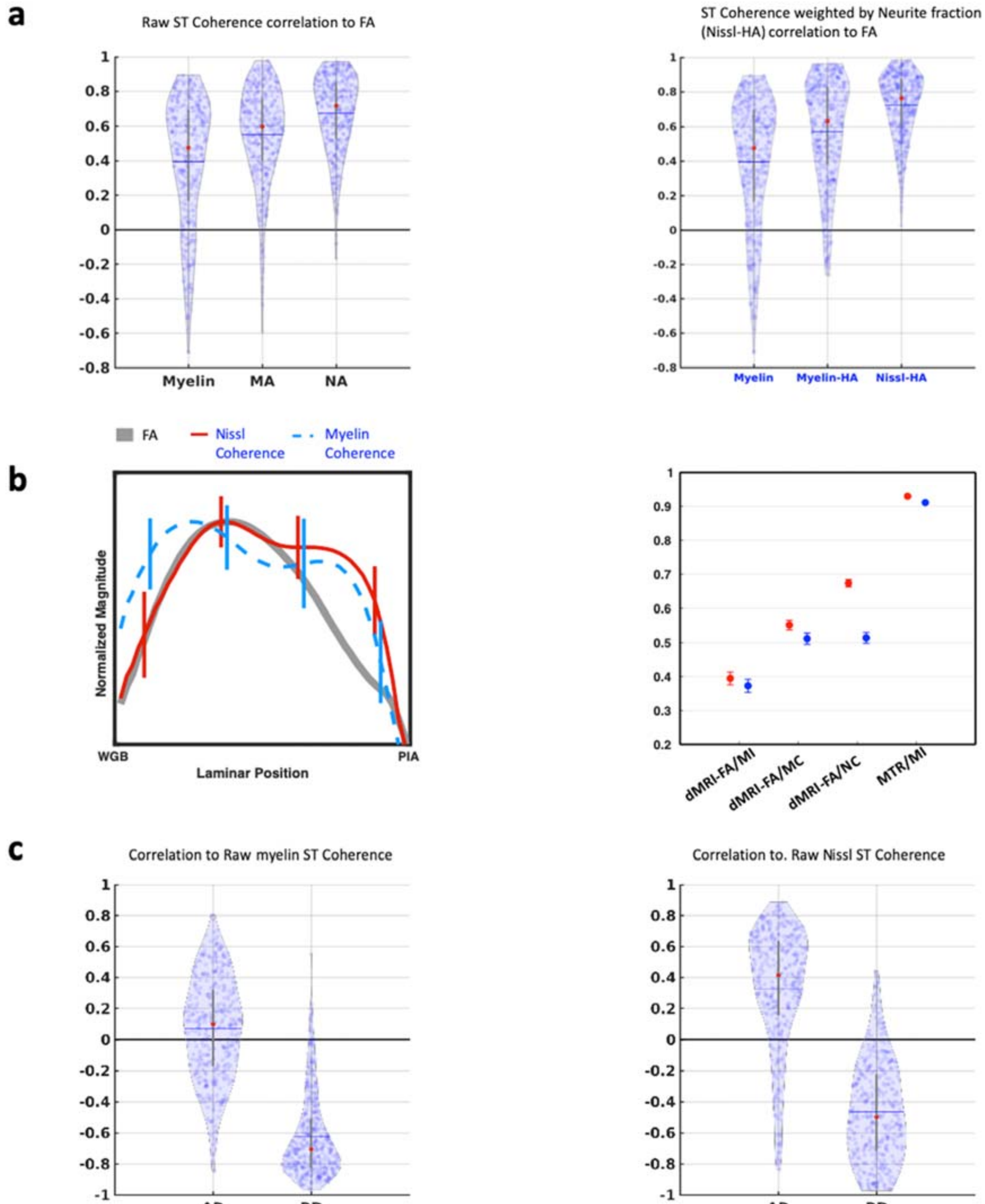


***SI Figure 5*** *Repeating the main results using raw Nissl and myelin ST Coherence rather than neurite density weighted Nissl-HA and myelin-HA.* ***a)*** *As figure 4a, correlation of histological raw histology ST coherence and myelin intensity to dMRI-FA.* ***b)*** *As Figure 5 Average laminar profiles of FA, Raw Nissl Coherence and myelin coherence across all parcels, and the results of a spatial randomization condition to test the regional specificity of parcel-wise histology to MRI correlations.* ***c)*** *As figure 6 correlations of Axial and Radial diffusivities to raw Nissl and myelin Coherence. The red circle shows the median value. The horizontal gray bars show the mean value, and the vertical gray bars show lower and upper quartiles.*

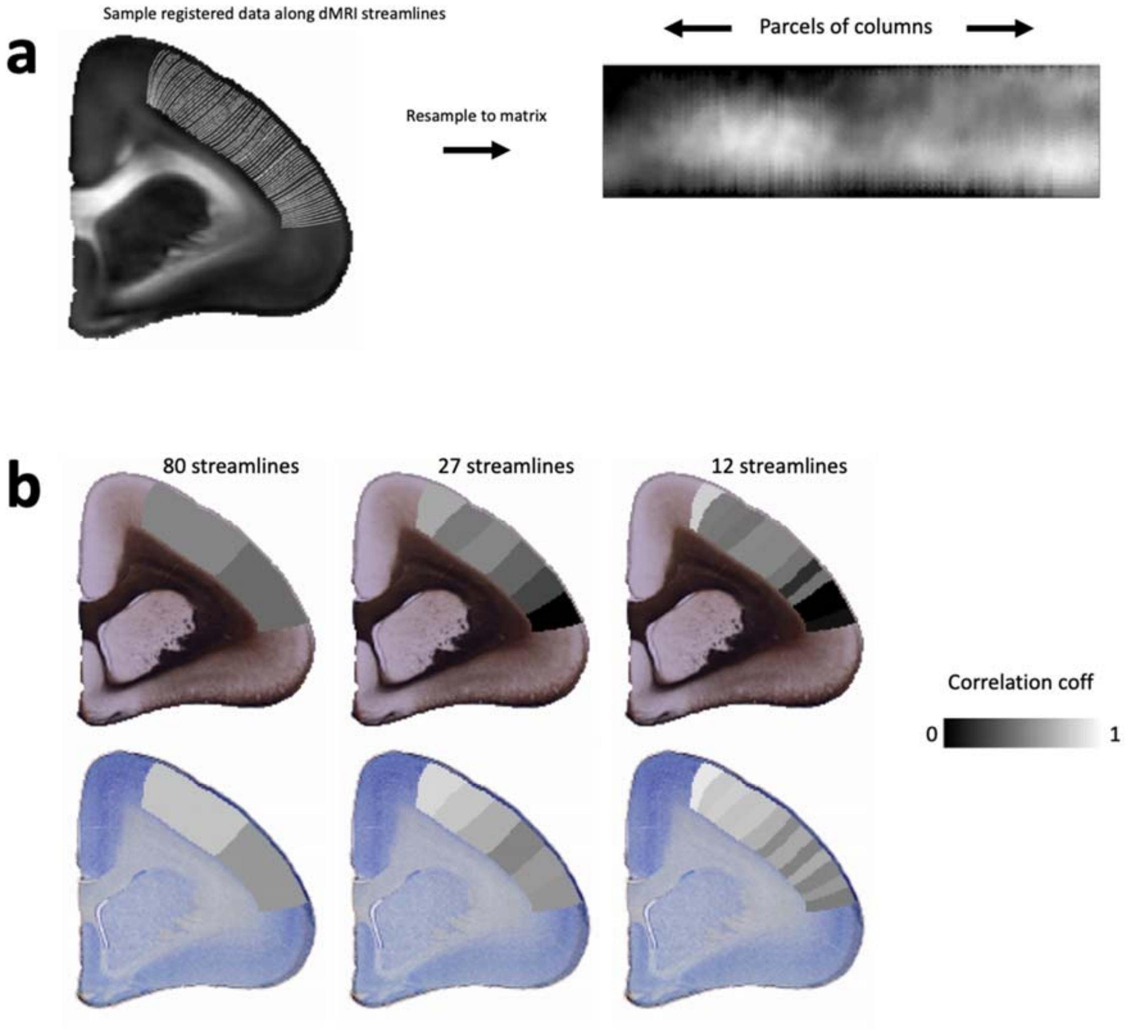


***SI Figure 6*** *Gray matter parcellation was achieved by sampling pixel values along streamlines and resampling into a matrix.* ***a)*** *pixels sampled along evenly spaced streamlines are resampled into a matrix for parcellation analysis* ***b)*** *correlation coefficients of parcels of 80, 27 and 12 streamlines displayed in image space.*

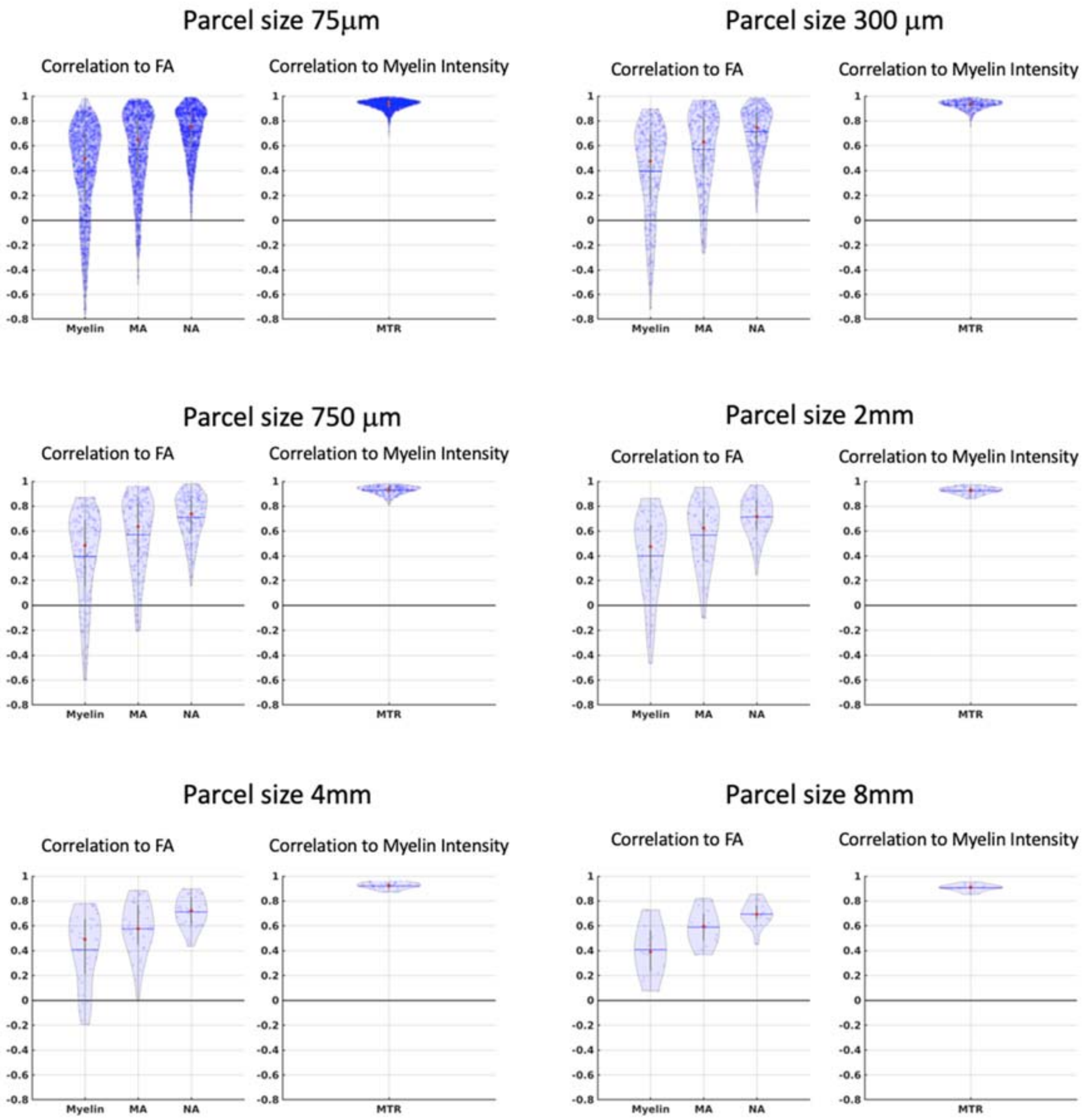


***SI Figure 7*** *Correlation of histology variables to dMRI-FA and MTR to myelin intensity, repeated with different cortical parcel sizes in the horizontal direction. Parcel size refers to the width of the parcel along the pial boundary. The mean parcel correlation is robust to parcel sizes ranging from 75μm to 8mm*

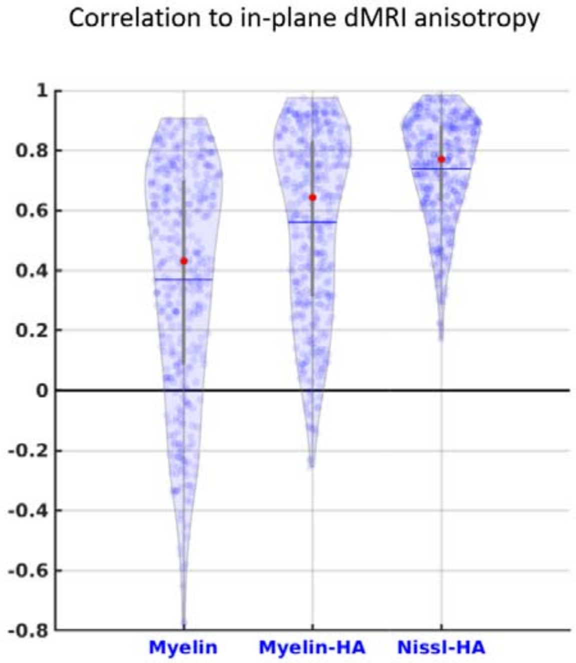


*SI Figure 8 Distribution of correlation coefficients between histology variables and in-plane dMRI anisotropy rather than dMRI-FA.*

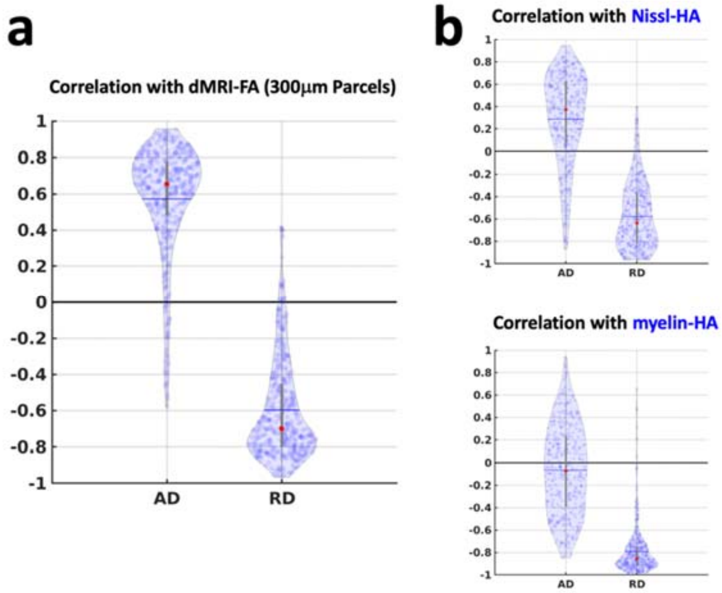


***SI Figure 9*** *Distribution of correlation coefficients between histological and dMRI anisotropy on one hand and directional diffusivities on the other. Nissl-HA exhibits a profile that is more similar to dMRI-FA than that of myelin-HA.* ***a)*** *dMRI-FA correlates to increases vertical diffusivity (AD) and decreases in horizontal diffusivity (RD).* ***Top Right*** *Nissl histological anisotropy (Nissl-HA) exhibits modest positive correlation to AD and negative correlation to RD, indicating that Nissl-HA and dMRI are associated with a similar mechanism of diffusion anisotropy.* ***Bottom Right*** *myelin histological anisotropy exhibits no mean correlation to vertical diffusivity and very strong correlation to horizontal diffusivity, as predicted from previous studies*

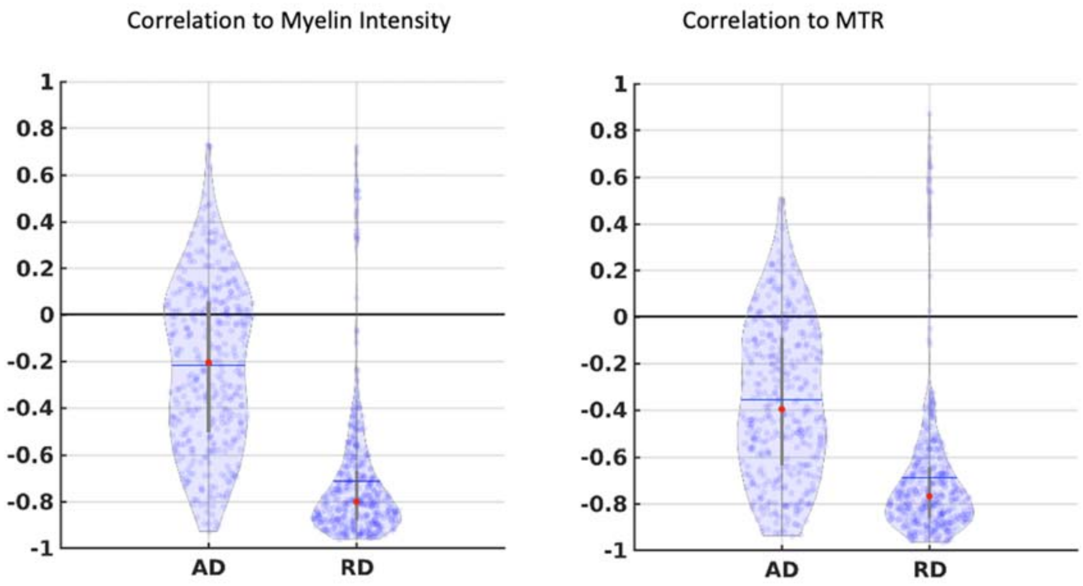


***SI Figure 10*** *Distribution of parcel correlation coefficients of axial and radial diffusivities to myelin intensity and MTR.*

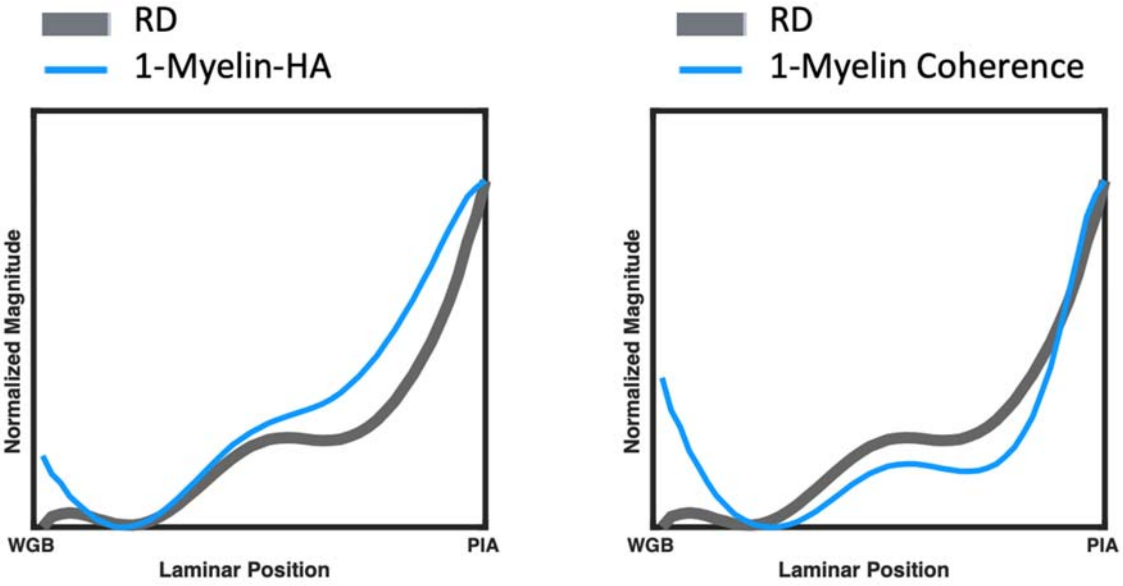


***SI Figure 11*** *Average laminar profiles of Radial diffusivity, inverse myelin-HA and inverse myelin ST coherence. Radial diffusivity closely covaries with the inverse of myelin anisotropy*

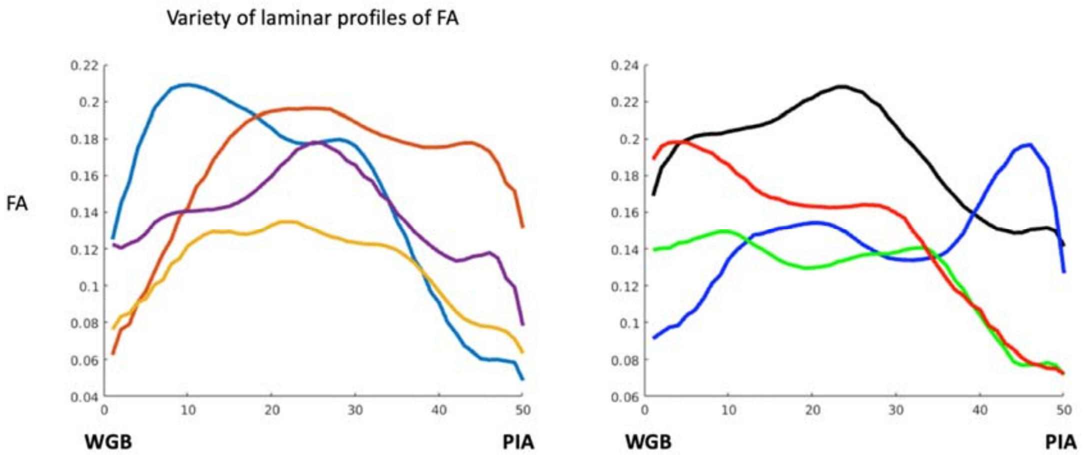


***SI Figure 12*** *Diverse trajectories of FA through the GM suggest different cortical areas may be discriminated*

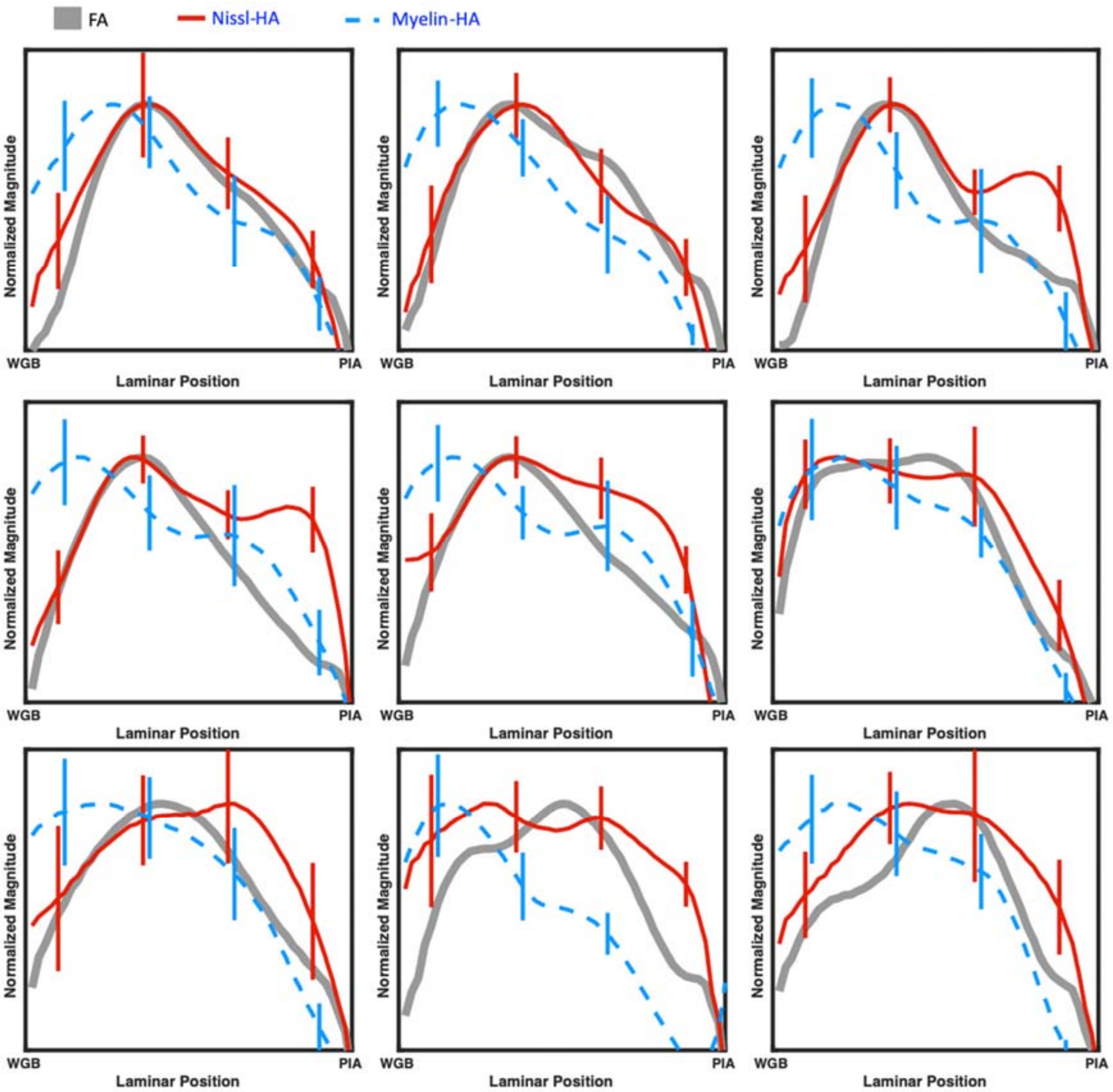


***SI Figure 13*** *Average laminar profiles of FA, Nissl-HA and myelin-HA in each of the nine non-linearly registered ROI*

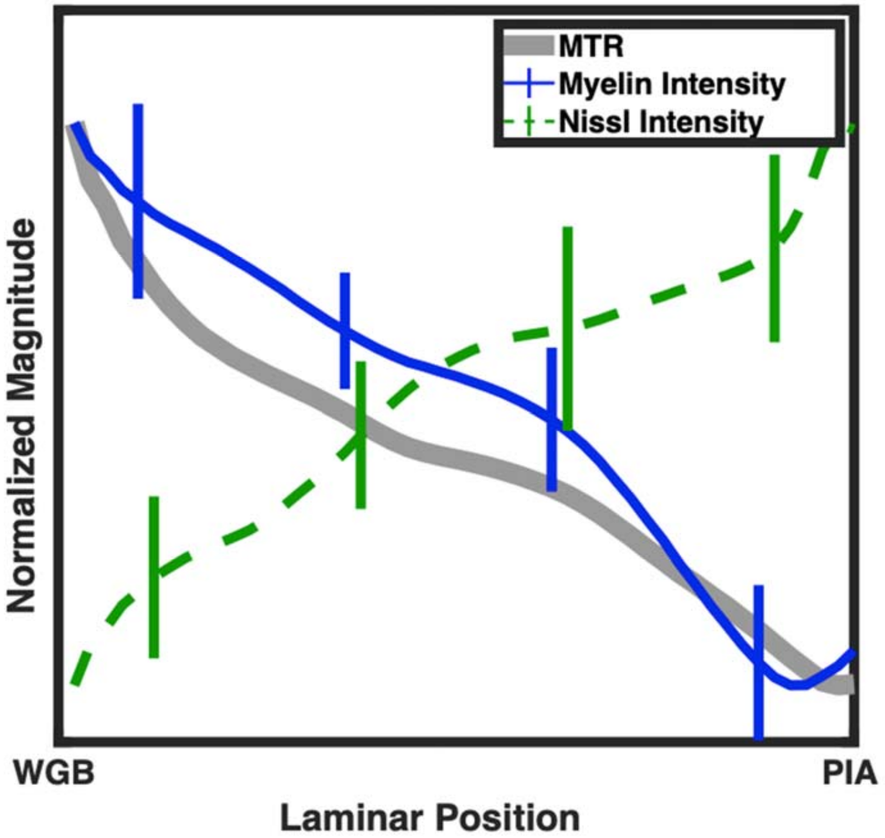


***SI Figure 14*** *Average laminar profiles of MTR, myelin stain intensity and Nissl stain intensity across all parcels,*

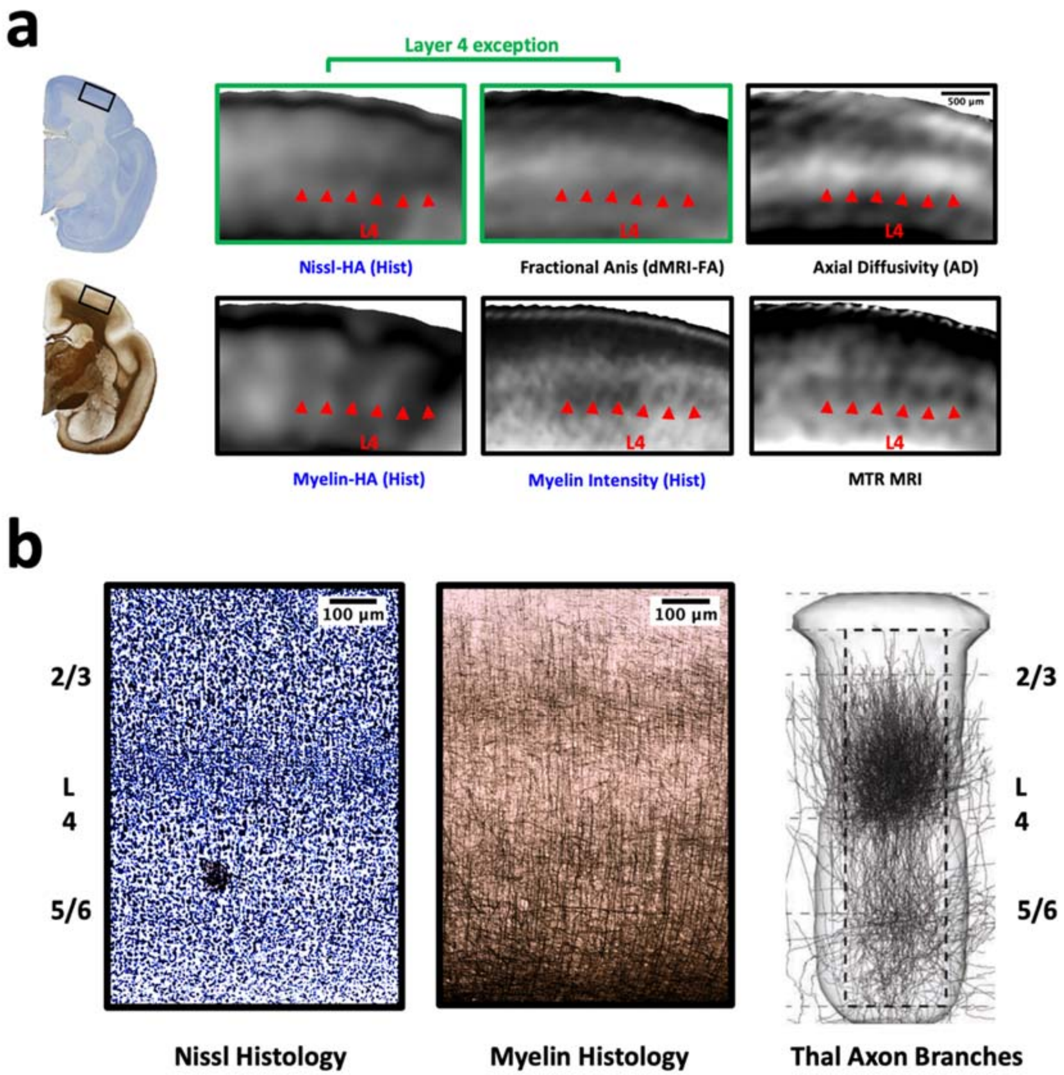


***SI Figure 15*** *In granular cortex, layer 4 is identifiable in Nissl histology by many small cell bodies and in myelin histology by a local decrease in stain intensity. In dMRI data layer 4 is associated with a local increase in Axial diffusivity (AD) and in MTR data by a band of low intensity.* ***a)*** *dMRI, MTR and histologically derived contrasts.* ***B) Left*** *Illustrative detail of the Nissl histology from an ROI of granular cortex. The internal granule layer is visible as a band of small, round cells.* ***Middle*** *the same region in myelin histology exhibits a gap in myelination. CLAHE contrast enhancement was applied uniformly across both myelin and Nissl images for display* ***Right*** *The highly branched, unmyelinated layer 4 terminals of axons from 8 thalamic afferent cells in rat Barrel cortex, reproduced from*

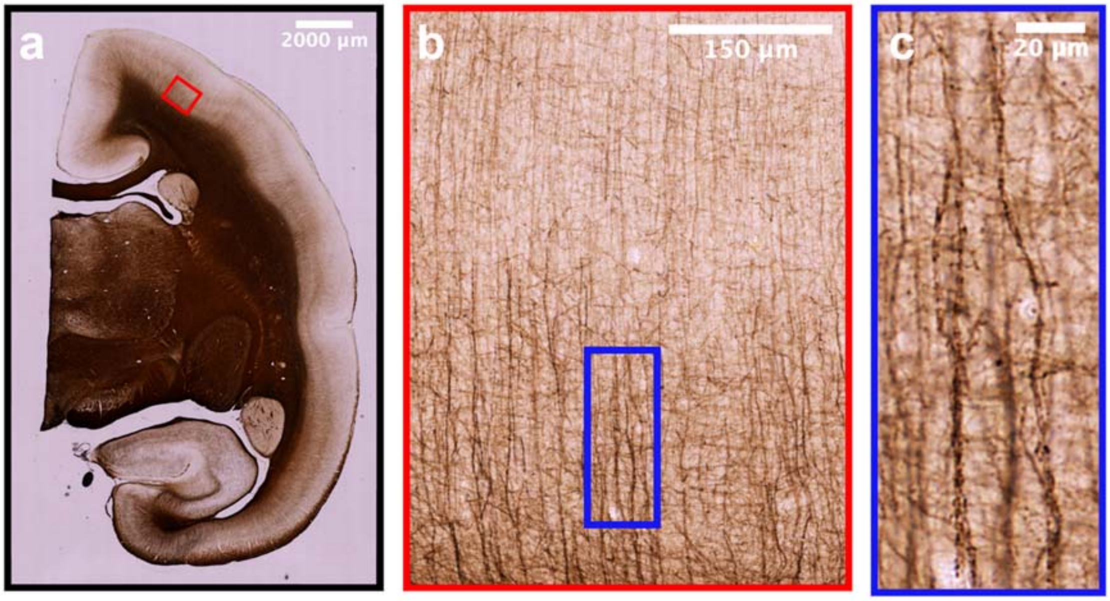


*SI Figure 16 individual myelinated axons can be discriminated, even in myelin rich deeper layers*

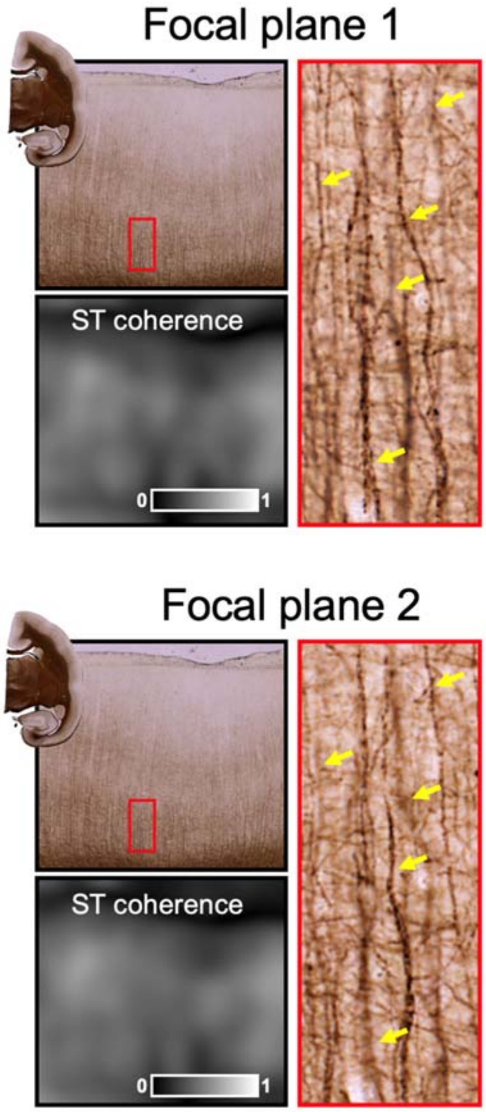


*SI Figure 17 Different myelinated fiber populations are resolved sharply using different focal depths within the 50µm thickness of the myelin stained section. However, the ST Coherence map over 150µm is quantitatively comparable, since the distribution of edge orientations is similar in both cases.*
